# An antigenic atlas of HIV-1 escape from broadly neutralizing antibodies

**DOI:** 10.1101/406355

**Authors:** Adam S. Dingens, Dana Arenz, Haidyn Weight, Julie Overbaugh, Jesse D. Bloom

## Abstract

Anti-HIV broadly neutralizing antibodies (bnAbs) have revealed vaccine targets on the virus’s Env protein and are themselves promising immunotherapeutics. The efficacy of bnAb-based therapies and vaccines depends in part on how readily the virus can escape neutralization. While structural studies can define contacts between bnAbs and Env, only functional studies can define mutations that confer escape. Here we map how all single amino-acid mutations to Env affect neutralization of HIV by nine bnAbs targeting five epitopes. For most bnAbs, mutations at only a small fraction of structurally defined contact sites mediated escape, and most escape occurred at sites that are near but do not directly contact the antibody. The mutations selected by two pooled bnAbs were similar to those expected from the combination of the bnAbs’ independent action. Overall, our mutation-level antigenic atlas provides a comprehensive dataset for understanding viral immune escape and refining therapies and vaccines.

## Introduction

Over the last decade, a burgeoning number of broadly neutralizing antibodies (bnAbs) have been isolated from HIV-infected humans. These antibodies target conserved regions of Env that are promising vaccine targets (Kwong and Mascola, 2018). Additionally, their broad neutralizing activity and potential to direct the killing of infected cells make bnAbs promising antiviral immunotherapeutic drugs for HIV prevention, therapy, and cure strategies (Margolis et al., 2017; Pegu et al., 2017).

However, bnAbs face a formidable foe. HIV Env’s exceptional evolutionary capacity allows the virus to stay one step ahead of bnAbs during infection, and resistance often arises when bnAbs are therapeutically administered to infected animals (Klein et al., 2012; Poignard et al., 1999; Shingai et al., 2013) or humans (Bar et al., 2016; Caskey et al., 2015, 2017; Lynch et al., 2015a; Scheid et al., 2016; Trkola et al., 2005). Thus, defining mutations that mediate viral escape is essential to optimizing and evaluating bnAb immunotherapies and vaccines.

While extensive efforts have gone into structurally characterizing bnAb epitopes via X-ray crystallography and cryo-electron microscopy (cryo-EM), structures on their own are insufficient to completely define the *functional epitope* (Cunningham and Wells, 1993; Kelley and O’Connell, 1993), defined here as sites where mutations affect antibody neutralization of replication-competent virus. Making individual mutations to Env and performing neutralization assays can provide information on the functional effect of specific mutations, but even the largest studies employing one-at-a-time mutagenesis can only assay a small fraction of all possible Env mutations.

We recently described mutational antigenic profiling, a massively parallel experimental approach to quantify how all single amino-acid mutations to Env affect antibody neutralization (Dingens et al., 2017). This approach involves generating libraries of HIV that carry all Env amino-acid mutations compatible with viral replication, incubating these libraries with or without an antibody, infecting T cells, and using deep sequencing to quantify the enrichment of each mutation in the selected versus non-selected libraries. Here, we apply this approach to a panel of nine bnAbs that target five Env epitopes, as well as a pool of two bnAbs. The resulting maps of viral escape provide comprehensive mutation-level views of the functional interfaces between HIV and bnAbs.

## Results

### Complete maps of viral escape from a panel of bnAbs

To gain a broad picture of viral escape, we selected nine bnAbs targeting the five best-characterized epitopes on Env (Figure 1A). Specifically, the bnAb panel includes the CD4 binding site (CD4bs) bnAbs VRC01 (Wu et al., 2010) and 3BNC117 (Scheid et al., 2011), the V3/N332 glycan supersite bnAbs PGT121 (Walker et al., 2011) and 10-1074 (Mouquet et al., 2012), the V2 glycan/trimer apex bnAbs PG9 (Walker et al., 2009) and PGT145 (Walker et al., 2011), the fusion peptide and gp120/gp41 interface bnAbs PGT151 (Falkowska et al., 2014) and N123-VRC34.01 (subsequently referred to as VRC34.01, Kong et al., 2016), and the membrane proximal external region (MPER) bnAb 10E8 (Huang et al., 2012). The binding footprints of these antibodies have been previously characterized using structural techniques (Figure 1B), allowing us to compare the *structural* epitopes with the *functional* epitopes defined by this study.

**Figure 1:**
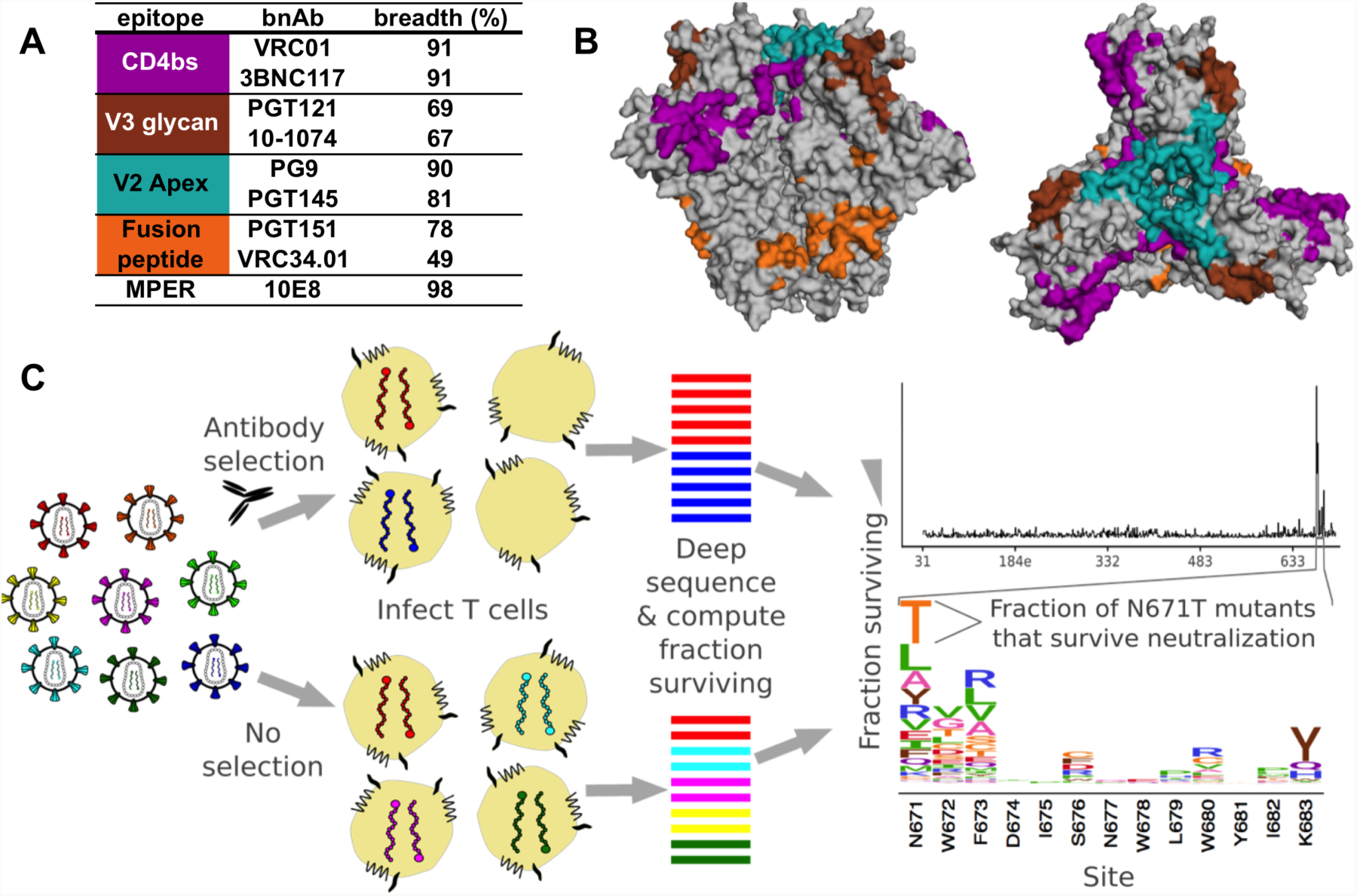
Schematic of mutational antigenic profiling of a panel of bnAbs. **A.** The bnAb panel. Breadth measures are from the 100 most commonly used viruses on LANL’s CATNAP (Yoon et al., 2015). **B.** For each epitope, structurally defined antibody contact sites are indicated by colors on the side and top view of the BG505 SOSIP Env trimer (PDB:5FYL). **C.** The mutational antigenic profiling experimental workflow, and example data from bnAb 10E8. A mutant virus library is incubated with or without and antibody before infecting SupT1.CCR5 cells. Non-integrated viral cDNA from infected cells is deep sequenced to quantify the frequency of each *env* mutation in both the antibody-selected and mock-selected conditions, and the overall fraction of the virus library that survives antibody neutralization is quantified via qPCR. The fraction of each mutant that survives neutralization is plotted at the site level in line plots, and at the mutation level in logoplots. The height of each letter is proportional the fraction of the virions with that amino acid that survived antibody selection in excess of the overall library average.

We mapped escape from these antibodies using the BG505.T332N Env, which is from a transmitted-founder subtype A HIV strain (Wu et al., 2006). This Env trimer is used widely in structural and vaccination studies (Sanders et al., 2013; Ward and Wilson, 2017). We used viral libraries that were previously generated by making all possible amino-acid mutations to the ectodomain and transmembrane domain of Env (Haddox et al., 2018). There are 19 amino-acid mutations × 670 sites = 12,730 such amino-acid mutations, and our libraries contain the subset of these mutations that is compatible with viral growth in cell culture.

To quantify how each of these mutations affect HIV’s antibody sensitivity, we neutralized independently generated mutant virus libraries at an ∼IC_95_-IC_99.9_ antibody concentration, and deep sequenced the *env* genes of viruses that were able to infect cells in the presence of antibody (Figure 1C). For each antibody we performed at least two replicates using independently generated viral libraries (Figure S1). We performed parallel control experiments without antibody, and compared the relative frequency of each Env mutation in the antibody-selected library to the non-selected control. By scaling this relative frequency by the overall fraction of the entire library that survived the antibody selection, we estimated the fraction of virions with each mutation that survive the selection, hereafter termed the *fraction surviving* (Doud et al., 2018). To highlight escape mutations, we plotted the excess fraction surviving above the overall library average.

Each antibody reproducibly selected mutations at just a small subset of Env sites (Figure 2A, Figure S1). The entire mutation-level maps of viral escape from each antibody are plotted across the entire mutagenized portion of *env* in File S1. Antibodies targeting the same epitope tended to select mutations in similar regions of Env, and these mutations cluster in three-dimensional structure in or near the antibody-binding footprint (Figure 2A). This is clearly exemplified by PG9 and PG145, where the selected positions largely overlapped. Note also that the effect size of mutations varied across antibodies (compare the y-axes in Figure 2A) as did the apparent noise in the plots; the implications of this are discussed in more detail in the final section of the Results.

**Figure 2:**
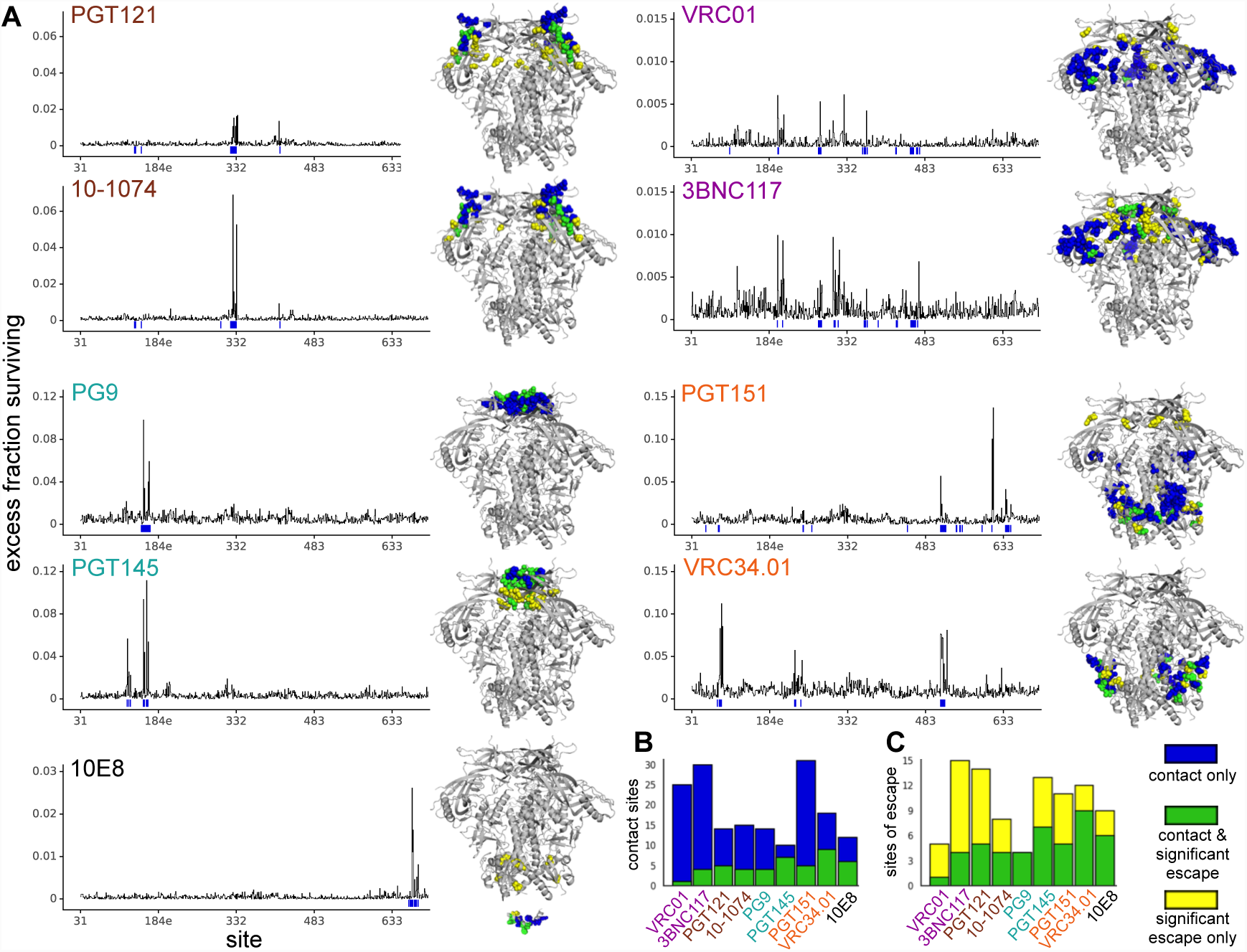
Env-wide escape profiles in relation to amino-acid positions that contact the bnAb. **A.** The line plots show the excess fraction surviving antibody neutralization averaged across all mutations at each site. Structurally defined contact sites are indicated by a blue line. The structures show the BG505 Env SOSIP trimer (PDB:5FYL) with sites of significant escape colored yellow, contact sites colored blue, and overlap between these sets of sites colored green. For 10E8, the MPER peptide structure (which is absent from the SOSIP trimer) is also shown (PDB:4G6F). **B.** Bars give the number of structurally defined contact sites for each antibody, with green indicating the contact sites that are also sites of significant escape. **C.** Bars give the number of sites of significant escape for each antibody, with green indicating the sites of escape that are also contact sites. Note that the green bars encompass the same sets of sites in panels B and C.

To rigorously compare the overlap between structural contacts and sites of viral escape, we identified sites of significant functional escape from the mutational antigenic profiling data, and sites of physical contact between the bnAb and Env from published structures. We defined significant sites of viral escape by fitting a gamma distribution to the measured antigenic effects of mutations at each site, and identified sites where the antigenic effects were larger than expected from this distribution at a false discovery rate of 0.01 (Figure S2). We defined sites of structural contact as Env residues that were within 4 Å of the antibody in available structural models, only considering non-hydrogen atoms (see Methods for details).

For most antibodies, only a small fraction of the structurally defined contact sites were also sites of significant viral escape (Figure 2B, Figure S3A). Further, we identified numerous sites of escape outside the structurally defined epitope for most antibodies. The extent to which escape occurred at sites that directly contact the antibody differs considerably across bnAbs, ranging from all significant sites of escape for PG9 to only one of five sites of escape for VRC01 (Figure 2C, Figure S3A). The sites of escape that do not directly contact the antibody are usually near the structurally defined epitope, in the 5-10 Å range (Figure S3A). However, a few sites of escape are more distant from the structurally defined epitope (Figure 2A, Figure S3B).

While our maps of escape include most mutations previously identified using individual BG505 point mutant pseudoviruses in TZM-bl neutralization assays (Kong et al., 2016; Lee et al., 2017), we also uncovered many previously uncharacterized sites of escape. We generated and tested BG505 point mutant pseudoviruses in TZM-bl neutralization assays for three antibodies, testing 16 to 19 point mutants for each antibody. The measurements from the mutational antigenic profiling were well correlated with the fold change in IC_50_ from TZM-bl neutralization assays for all antibodies tested (PGT121: R=0.76, n=16; 10-1074: R=0.81, n=16; VRC01: R=0.69, n=19) (Figure S4). In the next few subsections, we focus on each Env epitope individually.

### Escape from V3 glycan supersite bnAbs

The two anti-V3 antibodies PGT121 and 10-1074 are clonal variants that arose in the same infected individual (Mouquet et al., 2012). However, there were intriguing differences between the two antibodies in the specific mutations that mediated escape in our experiments, as well as the overall effect sizes of escape mutations (Figure 3A, 3B, note the different y-axis scales in the two panels). For instance, mutations to site 325 had a larger effect on 10-1074 than PGT121, whereas mutations at site 327 had similar effects. We validated the differential effects of mutations to site 325 by testing three different mutants at this site in TZM-bl neutralization assays: the maximal effect for 10-1074 was a 27-fold increase in IC_50_, while the maximal effect for PGT121 was just a 1.7-fold increase (D325E; Figure S4). Our mutational antigenic profiling also indicated that mutations that eliminated the N332 glycan had a larger effect for 10-1074 than PGT121 (Figure 3A,B), consistent with a prior study that examined binding to gp120 (Mouquet et al., 2012). In contrast, at most other sites in the epitope (such as 323, 327, and 330), the overall effects of mutations were similar between the two antibodies (Figure 3A,B).

**Figure 3:**
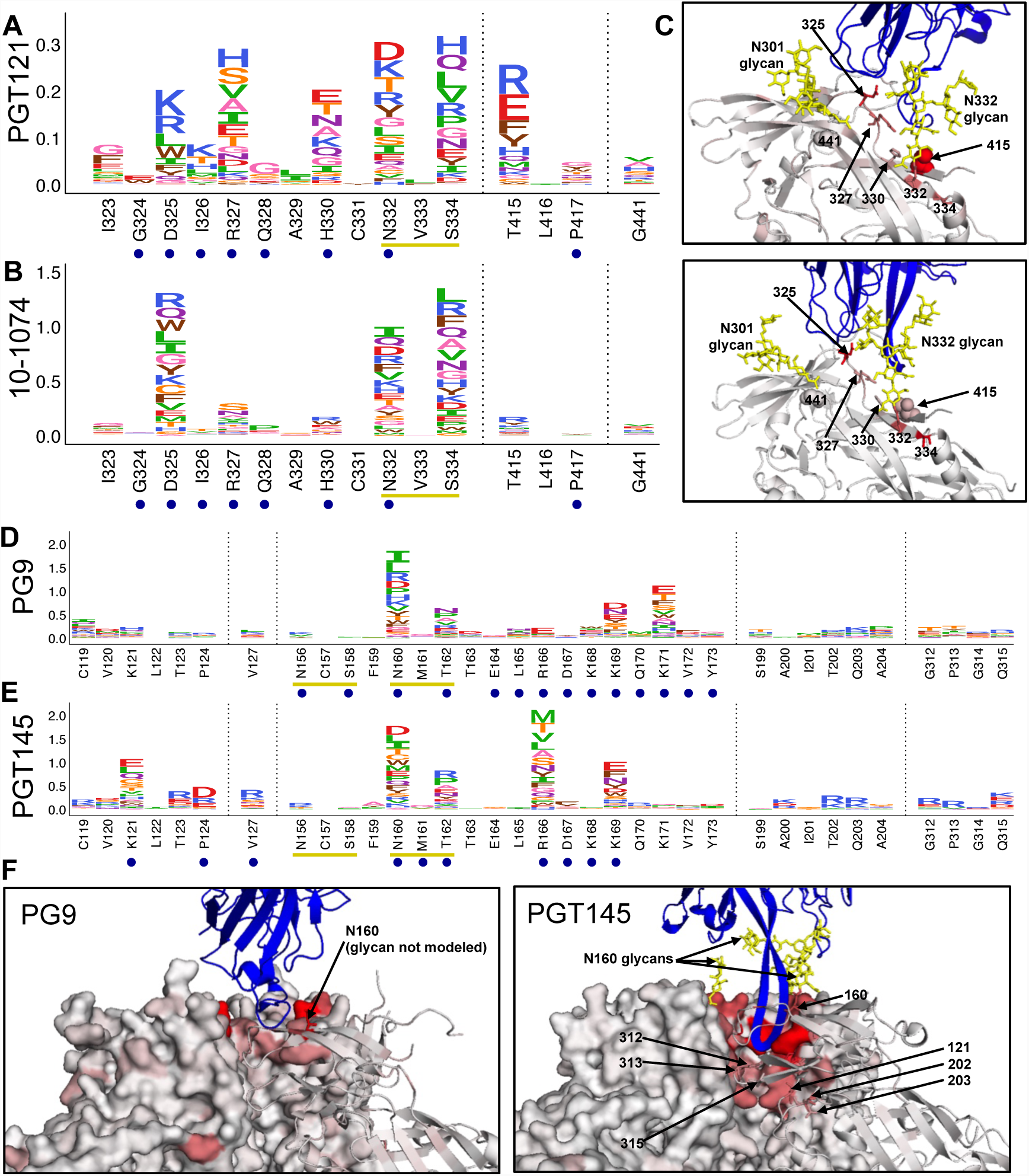
Escape From V3 glycan supersite and V2 apex bnAbs. **A**, **B.** Escape profiles for V3 glycan supersite bnAbs PGT121 and 10-1074. Letter heights indicate the excess fraction surviving for each mutation. Blue circles indicate structurally defined contact sites, and yellow underlines indicate a N-linked glycosylation motif. Logoplots that show escape across Env are in File S1. **C.** V3 glycan supersite antibodies are shown in blue, and Env is colored according to the maximum excess fraction surviving at each site. Note that for PGT121, the closely related clonal variant PGT122 structure is used in lieu of a PGT121 structure (PDBs: 5FYL and 5T3Z respectively). **D**, **E.** Escape profiles of V2 glycan/apex antibodies PG9 and PGT145, presented in the same manner as A, B. **F.** V2 glycan/apex antibodies are shown in blue, and Env is colored according to the maximum excess fraction surviving at each site (PDBs: 5VJ6 and 5V8L respectively).

There are also differences in *which* mutations at a given site escape each antibody. For example, while the overall effect of all mutations at site 330 is similar between 10-1074 and PGT121, H330R escapes 10-1074 but not PGT121 (Figure 3A,B, validated by TZM-bl neutralization assays in Figure S4). Even among small-effect mutations, TZM-bl neutralization assays validated the results of mutational antigenic profiling. For example, mutations at site 325 had disparate effects: D325E had a large effect on 10-1074 but a negligible effect PGT121, D325S had a small effect for 10-1074 but no effect for PGT121, and D325N had no effect on either antibody (Figure 3A,B and validated in Figure S4).

Many aspects of our maps of escape are consistent with prior knowledge about the epitopes of these two antibodies (Garces et al., 2014; Mouquet et al., 2012; Sok et al., 2014, 2016). For example, eliminating the targeted N332 glycan via mutations to N332 and S334 resulted in escape from both antibodies, as did antibody-specific mutations in the ^324^GDIR^327^ motif (Figure 3), a conserved region of this epitope that is involved with CCR5 co-receptor binding (Sok et al., 2016).

However, we also identified escape at sites not previously implicated as being part of the functional epitope. For instance, viral escape from both antibodies occurred via mutations at site 415 in V4, and to a modest but reproducible extent, site 441 (Figure 3A, 3B). We validated that mutations at each of these sites resulted in escape from both antibodies in TZM-bl neutralization assays (Figure S4). Neither of these sites directly interact with either antibody; site 415 it is close to other structural contacts as well as the N332 glycan, and site 441 in the β22 strand neighbors the N301 glycan.

### Escape from V2 apex bnAbs

For the V2 apex antibodies PG9 and PGT145, escape occurred via eliminating the N160 glycan at the heart of the epitope (Figure 2A, 3D, 3E). Additionally, escape occurred at structurally defined contact sites at the trimer apex for PG9, and at the trimer apex and interface for PGT145. For both antibodies, prior studies suggest that binding is driven by electrostatic interactions with positively charged Env residues (Lee et al., 2017; McLellan et al., 2011; Wang et al., 2017). Mutations to these sites resulted in viral escape (including residues R166, K169, and K171 for PG9, and K121, R161, K169 for PGT145), with charge swaps often resulting in the greatest extent of escape (Figure 3D, 3E). Structural studies indicate that the long HCDR3 arm of PGT145 reaches into the trimer interface; existence of this epitope has been hypothesized to result from a balancing act of a “push” from inter-protomer charge repulsions at the trimer interface and a “pull” of hydrophobic interactions between variable loops across protomers at the trimer apex (Lee et al., 2017).

While escape from PGT145 occurred via eliminating the epitope’s positive charges, escape also occurred via *introducing* charges at sites where the wildtype residue is not charged. These included sites 123, 124, and 127 in or very near the epitope, as well as more distant sites encircling the epitope, including sites 200, 202, 203 in the β3-β4 loop, and 312, 313, and 315 at the tip of the V3 loop (Figure 3F). These mutations presumably also affected the charge repulsions at the trimer interface and/or overall trimer conformation, disrupting the electrostatic balancing act that is crucial for PGT145 binding.

### Escape from CD4bs bnAbs

Escape from CD4bs bnAbs VRC01 and 3BNC117 occurred in both the canonically defined CD4bs epitope and other sites distal to the CD4 binding site (Figure 2A, Figure 4). In the CD4 binding site, mutations to site 279 in loop D and site 369 in the CD4 binding loop escaped both antibodies (Figure 4A). With the exception of sites 279 and 280, the specific amino-acid mutations in loop D that mediated escape differed between VRC01 and 3BNC117.

**Figure 4:**
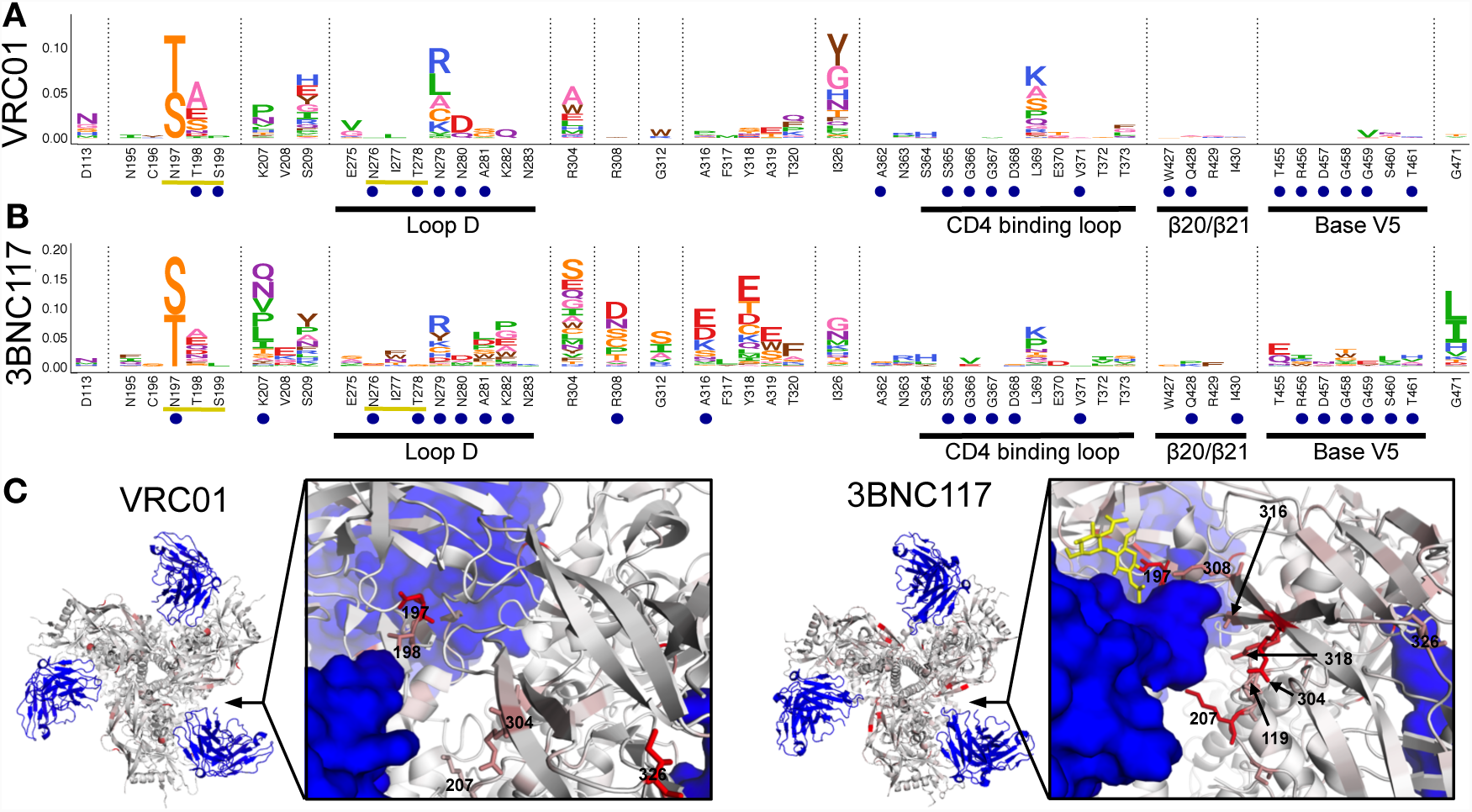
Escape from CD4bs bnAbs. **A**,**B.** Escape profiles for CD4bs bnAbs VRC01 and 3BNC117. Letter heights indicate the excess fraction surviving for each mutation. Blue circles indicate structurally defined contact sites, and yellow underlines indicate a N-linked glycosylation motif. Portions of the canonical CD4bs epitope are underlined in black and labeled. Logoplots that show escape across Env are in File S1. **C.** Antibodies are shown in blue, and Env is colored according to the average fraction surviving at each site (PDBs: 5FYK and 5V8M respectively).

The largest-effect mutations for both antibodies introduced a serine or threonine in place of the asparagine at site 197. The N197 glycan is part of a glycan fence that shields the CD4bs (Crooks et al., 2017). N197S/T both eliminate this N197 glycan and introduce a new potential N-linked glycosylation motif (PNG) at N195. Since escape was only mediated by S/T at site 197, these data suggest that eliminating N197 alone does not result in viral escape, but shifting the N197 glycan to N195 does. We validated these observations using point mutants in TZM-bl neutralization assays: simply eliminating the N197 PNG via N197E resulted in ∼50 fold *more* potent neutralization by VRC01, while N197S resulted in viral escape, increasing the IC_50_ by 27-fold relative to wildtype (Figure S4).

Escape from both antibodies also occurred via D113N, which introduces a PNG at site 113. Site 113 is in the trimer interface distant from the CD4bs (Figure 4C), suggesting this mutation may affect exposure of the CD4bs epitope by altering trimer conformation or dynamics. We validated that D113N resulted in escape from VRC01 using a TZM-bl neutralization assay (Figure S4). While it is unknown if the PNG created by D113N is indeed glycosylated, these data show that altering potential glycosylation sites both near and distal to the epitope can affect CD4bs bnAb neutralization.

Escape from 3BNC117 also occurred at numerous sites near where the antibody’s HCDR3 arm makes inter-protomer contacts, including sites in V3 (sites 304, 308, 312, 316-320), and at the base of the β3-β4 loop (sites 207, 209) (Figure 4B). This quaternary nature of the 3BNC117 epitope was first postulated based on early trimer structures (Lyumkis et al., 2013), and higher resolution cryo-EM of BG505 trimer in complex with 3BNC117 (Lee et al., 2017) confirmed that 3BNC117 directly interacts with residues 207, 308, and 316 from the neighboring protomer (Figure 4B, 4C). It has been previously reported that mutations to site 207 result in decrease 3BNC117 binding (Liu et al., 2017). We also observed viral escape from 3BNC117 at site I326, a site distal to 3BNC117 near the base of the V3 loop that takes part in variable loop hydrophobic interactions that may regulate trimer dynamics (Lee et al., 2017).

Strikingly, while VRC01 does not make similar inter-protomer structural contacts as 3BNC117 (Stewart-Jones et al., 2016), we still observed escape at sites 207, 209, 304 and 326 (Figure 4A, 4C). We validated that I326Y results in escape from VRC01 (Figure S4), but has little effect on the V3-specific bnAbs 10-1074 and PGT121, despite these antibodies directly contacting this site.

### Escape from fusion peptide bnAbs

Maps of escape from PGT151 and VRC34.01 highlight the complex nature of the conformational fusion peptide and gp120/gp41 interface epitope (Figure 2A, 5A, 5B). Here, we reanalyzed VRC34.01 mutational antigenic profiling data from a prior study (Dingens et al., 2018) quantifying the effects of mutations using the *fraction surviving* metric rather than the *differential selection* metric used in the earlier study, and compared these data to the BG505 Env escape from PGT151 reported here. While both antibodies contact the 6 N-terminal residues of the fusion peptide (512-517), escape from PGT151 is focused on just the 3 N-terminal residues of this peptide (512-514), while escape from VRC34.01 is mediated by numerous mutations to sites 512-516 and 518. The structural footprints of both antibodies center on the fusion peptide, but they contact distinct glycans and protein regions of gp120 and gp41. Again, their *functional epitopes* include distinct subsets of these of protein residues and glycans (Figure 5A, 5B). For both antibodies, there are also numerous sites of significant escape at non-contact sites near the epitope (Figure 5A, 5B). For PGT151, we also identified sites of escape at more distant residues in V3; the mechanisms of escape at these sites are unclear.

**Figure 5:**
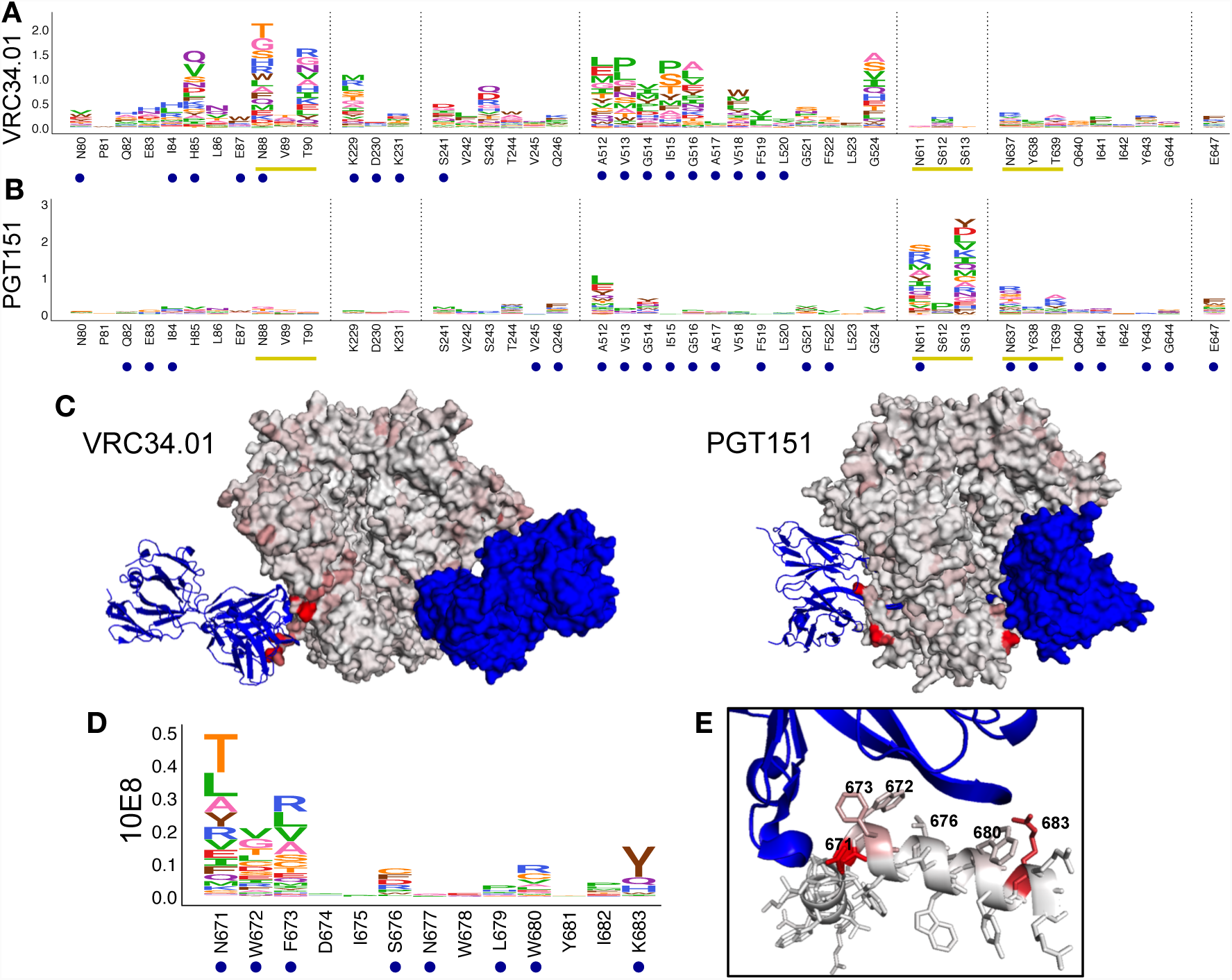
Escape from fusion peptide and MPER bnAbs. **A**, **B.** Escape profiles for fusion peptide bnAbs VRC34.01 and PGT151. Letter heights indicate the excess fraction surviving for each mutation. Logoplots that show escape across Env are in File S1. **C.** Fusion peptide antibodies are shown in blue, and Env is colored according to the maximum excess fraction surviving at each site (PDBs: 5I8H and 5FUU respectively). **D.** Escape profile for MPER bnAb 10E8, presented in the same manner as A, B. **E.** 10E8 is shown in blue, and the MPER peptide is colored according to the maximum excess fraction surviving at each site (PDB 4G6F).

### Escape from an MPER bnAB

Escape from the MPER-directed antibody 10E8 occurred predominantly in the structurally defined contact sites, with sites of escape localizing to one side of the MPER peptide alpha helix apical to 10E8 (Figure 5D, 5E) (Huang et al., 2012). This agrees precisely with prior studies (Huang et al., 2012; Kim et al., 2014). However, we identified two additional modest but significant sites of escape outside of the MPER peptide, at sites 609 and 643 in the C-C loop and HR2 domain of gp41 respectively (Figure 2A, Figure S2). Mutations at these sites may alter fusion kinetics and/or the presentation of the MPER epitope.

### Comparing our mutational maps with in vivo escape during bnAb immunotherapies in humans

Several of the bnAbs that we characterized have been used in human immunotherapy studies. Some of the escape mutations identified in our work overlap with mutations that arose in the humans during these studies (Table S1). For example, when 10-1074 was administered to HIV-infected individuals, viral escape mutations emerged at site 325 and the PNG that encompasses sites 332 and 334 (Caskey et al., 2017). These are the same three sites where the strongest selection is observed in our 10-1074 mutational antigenic profiling (Figure 3B, Table S1).

There is also considerable overlap between sites of escape we map *in vitro* and those that occurred *in vivo* during treatment of infected individuals with the CD4bs antibodies 3BNC117 or VRC01. For 3BNC117, the sites over overlap between our maps and human trials (Caskey et al., 2015; Schoofs et al., 2016) includes sites 182, 209, 279, 308, 318, and 471 (Table S1). Site 279 is one of the strongest sites of escape from 3BNC117 and VRC01 in our experiments. A mutation to site 279 was part of the viral escape pathway within the patient from whom VRC01 was isolated (Lynch et al., 2015b), and arose during VRC01 immunotherapy post treatment interruption (Bar et al., 2016). Mutations to site 279 also played a role in escaping a CD4bs targeted response in another patient (Wibmer et al., 2013) and during 3BNC117 immunotherapy of infected individuals (Caskey et al., 2015; Schoofs et al., 2016).

Intriguingly, our data may also be useful for identifying previously unappreciated escape mutations during immunotherapy. For example, after patient V10 underwent therapy with VRC01 (Bar et al., 2016) a rare amino acid variant at site 326 was fixed in the viral population (Table S1), but the potential significance of this mutation was not noted in the original publication since it is far from the structural epitope. Our mutational antigenic profiling shows that mutations at site 326 increase resistance to VRC01 (Figure 4A), demonstrating how comprehensive maps of mutational escape can aid in interpreting clinical data.

### Escape from pooled antibodies

Many immunotherapy studies are beginning to treat patients with combinations of bnAbs. For instance, one rapidly advancing set of clinical trials involves treating patients with equal concentrations of 3BNC117 and 10-1074 (NIH ClinicalTrials.gov Identifiers: NCT03526848, NCT02824536, NCT02825797). We therefore investigated how escape from a mix of these two antibodies compares to escape from each antibody individually.

We pooled the antibodies at equal concentrations, and then selected our viral libraries with the antibody pool (Figure 6, Figure S5). Escape from the pooled antibodies appeared to be a combination of the escape profiles from each antibody in isolation (Figure 6A, 6B). For example, we observed escape at sites 325, 332, and 334, likely associated with escape from 10-1074, as well as escape at sites 304, 308, and 471, which presumably affect 3BNC117 resistance.

**Figure 6:**
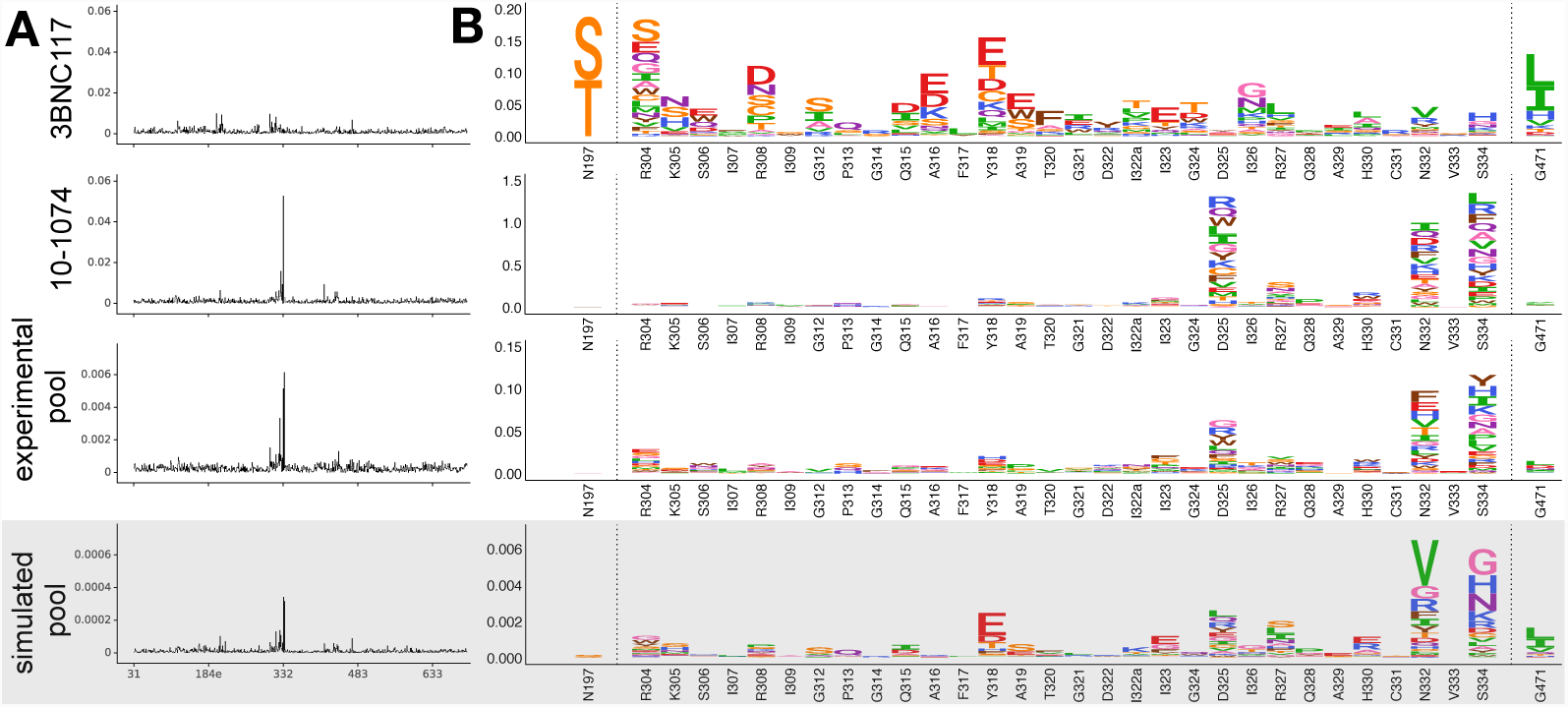
Escape from a 3BNC117 and 10-1074 pooled bnAbs. **A.** The excess fraction surviving neutralization averaged across all mutations at each site. Data from Figure 3 (10-1074) and Figure 4 (3BNC117) are re-plotted for relevant sites. For the pooled 3BNC117 and 10-1074 data, the mean value across six replicates is plotted. The simulated data is the product of each antibody’s mean excess mutation fraction surviving values. **B.** A logoplot zooming in on epitope regions for each dataset. In **A** and **B**, the simulated data is distinguished from the experimental data with a light grey overlay.

Importantly, escape from the pooled antibodies does not occur at sites where the escape mutation from one antibody sensitizes the virus to the other. For instance, the strongest escape mutations for 3BNC117 alone are N197S and N197T, which shift the N197 glycan to N195 (Figure 4B and Figure 6A). However, eliminating the N197 glycan *increases* the virus’s susceptibility to neutralization by antibodies targeting the same epitope as 10-1074 (Liang et al., 2016; Townsley et al., 2016). Mutations at site N197 are not selected by the pool of antibodies in our mutational antigenic profiling, presumably because any benefit with respect to escaping 3BNC117 is canceled out by increased susceptibility to 10-1074. This example demonstrates the potential for suppressing viral escape mutations by selecting antibodies with synergistic effects at specific sites.

To model the synergistic effects of antibodies on suppressing viral escape, we calculated the expected escape profile from a pool of 10-1074 and 3BNC117 by simply taking the product of each mutation’s excess fraction surviving value for each antibody (Figure 6). The rationale behind this calculation is that the expected fraction of virions with a mutation that should survive both antibodies is simply the product of the fraction that would survive each antibody individually. The escape profile predicted by this simple model closely matches the actual selection from the pooled antibodies (Figure 6).

Overall, these data suggest that no single amino-acid mutations robustly escape both 3BNC117 and 10-1074. Rather, the low level escape from the pooled antibodies appears to represent mutations that escape one antibody but have little effect on the other. Furthermore, the similarity of the experimentally measured escape profile for the pooled antibodies and the profile predicted from the product of the individual antibody profiles suggest that our maps of escape from single antibodies could be useful for computationally predicting the potential for escape from antibody pools.

### Quantifying the ability of the BG505 Env to escape each bnAb with single mutations

Anti-HIV bnAbs are often evaluated in terms of their breadth and potency against panels of naturally occurring viral strains. Our data offer the opportunity to calculate an alternative measure relevant to the potential efficacy of bnAb immunotherapies: the ability of single amino-acid mutations to increase the antibody resistance of a particular viral strain.

We used our mutational antigenic profiling to assess the ease of single-mutation escape of the BG505 Env from each antibody. First, we simply qualitatively examined the 100 largest effect-size mutations for each antibody (Figure 7A). For some antibodies (such as VRC34.01), there are many individual mutations that efficiently escape neutralization—but for other antibodies (such as VRC01), only a few mutations affect escape, and do so with only moderate size effects (Figure 7A). Interestingly, for all of the antibodies, at least some of the largest effect-size mutations are accessible by single nucleotide mutations, indicating that the genetic code only has moderate affects on the accessibility of escape mutations (Figure 7A).

**Figure 7:**
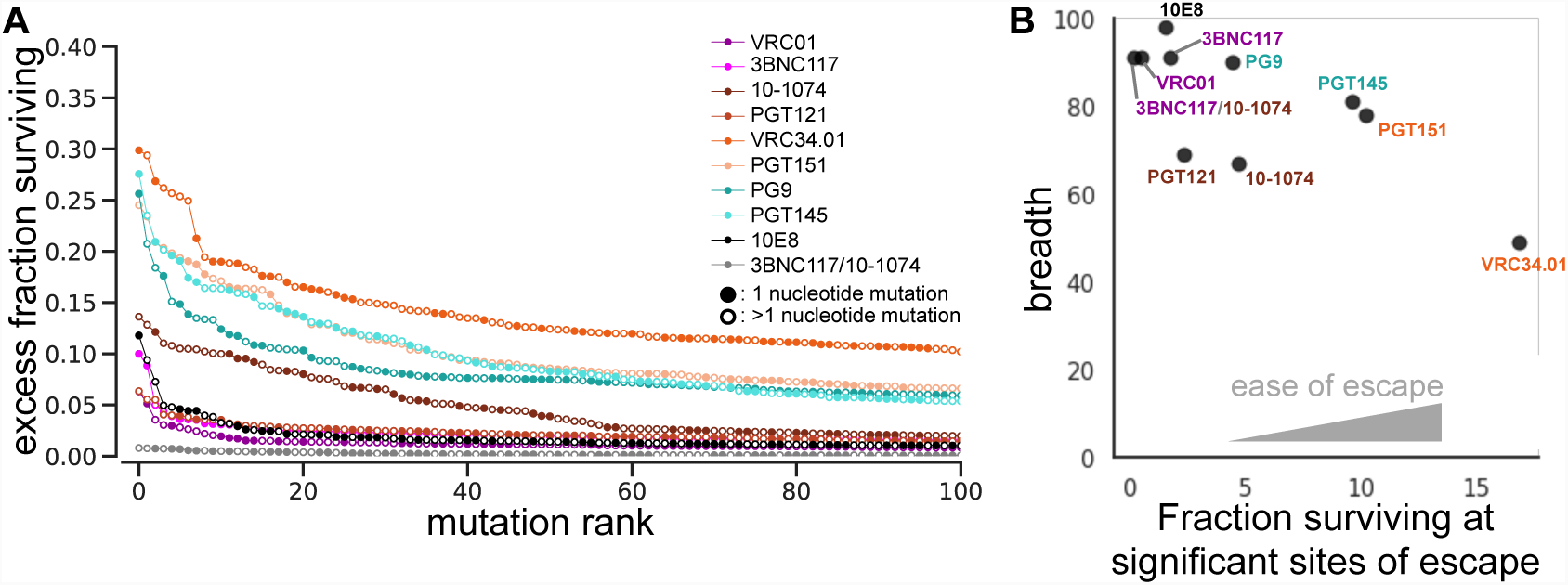
Quantifying the ability to escape each bnAb. **A**. The excess fraction surviving for the 100 largest effect mutations for each antibody. Closed circles indicate mutations that are one nucleotide mutation away from BG505 wildtype, while open circles indicate mutations that are 2 or 3 nucleotide mutations away from BG505 wildtype sequence. **B.** Each antibody’s breadth (as quantified in Figure 2) is plotted against the sum of the excess mutation fraction surviving values at all significant sites of viral escape.

To quantify the ease with which the BG505 Env can escape each antibody by single mutations, we summed the excess mutation fraction surviving values at each significant site of viral escape. This single-mutation ease of escape metric is moderately correlated with antibody’s breadth (Figure 7B). But especially for the broadest antibodies, there are differences between neutralization breadth on natural strains and ease of single-amino acid escape by BG505 (Figure 7B). For example, 10E8 has 98% breadth and VRC01 has 91% breadth, but BG505 has more capacity to escape 10E8 by single amino-acid mutations. These results highlight that, similar to influenza antibodies (Doud et al., 2018), HIV bnAb breadth on natural strains and the potential for single-mutation escape by any given viral variant are distinct measures, both of which may be useful for assessing the potential for viral antibody escape in clinical settings.

## Discussion

We have mapped how all single amino-acid mutations to the BG505 Env affect neutralization of replication-competent HIV by nine prototypical bnAbs. These maps of viral escape define the functional epitopes of these antibodies, which we show here are distinct from the structurally defined epitopes. For all antibodies, viral escape occurred at only a fraction of structurally defined contact sites, and many escape mutations occurred at sites outside the direct structurally defined epitope. This escape at non-contact sites often clustered close to the structural contacts, suggesting that altering a network of interacting sites near the structural epitope can disrupt antibody binding or neutralization. In a few cases, escape also occurred at sites more distant from the epitope, likely due to alterations in the conformation or dynamics of Env.

Others have previously noted that not all structurally defined contact sites affect antibody binding (Falkowska et al., 2012; Kelley and O’Connell, 1993; Li et al., 2011), and mutations outside of the structural epitope can affect antibody sensitivity (Back et al., 1993; Blish et al., 2008; Boyd et al., 2015; Bradley et al., 2016; O’Rourke et al., 2012; Sethi et al., 2013). However, our complete maps of escape mutations make it possible to systematically quantify the overlap between the functional and structural epitopes of bnAbs. Our work also highlights how incompletely prior structural and virological assays have defined the functionally relevant interactions between HIV and antibodies. Even though our study focused on some of the best-characterized anti-HIV bnAbs, we uncovered numerous sites in Env (often outside of the structural epitope) where mutations were not previously known to affect bnAb sensitivity.

One application for which our maps of viral escape may be of immediate use is evaluating bnAb immunotherapies. During immunotherapy trials, many viral mutations arise, and it is important to know which ones alter sensitivity to the bnAb used in the trial (Bar et al., 2016; Caskey et al., 2015, 2017; Lynch et al., 2015a; Scheid et al., 2016; Schoofs et al., 2016). While our maps of course do not perfectly predict escape that occurs *in vivo*, which could be influenced by many stochastic factors including the specific variant that infected the individual, they do define functional epitopes that can be used to help assess the potential antigenic significance of viral mutations. For instance, our map of the functional epitope of VRC01 allowed us to re-interpret the potential significance of a previously unremarked upon mutation outside the structural epitope that occurred during VRC01 immunotherapy.

We also examined how our maps of escape from individual antibodies compared to those generated using a pool of antibodies—an important question since combinations of bnAbs are a clear future direction in antibody immunotherapy (Wagh et al., 2016, 2018). We mapped escape from one such clinically relevant pool of two bnAbs, 3BNC117 and 10-1074. We found that there were no mutations that robustly escaped both antibodies. Further, the combined selection of the two antibodies was similar to that predicted from the product of the individual escape profiles. Therefore, antigenic maps for individual antibodies may be useful for modeling viral escape from combinations of antibodies.

Our data also provide a way to quantify the ease with which the BG505 Env can escape from each antibody via single mutations. While this ease of escape metric contains has many caveats specific to our experimental system (it only looks at single mutations to BG505 that support viral replication in cell culture), caveats also apply to other methods for estimating how likely a virus is to escape an antibody. For example, characterizing viral escape during immunotherapy in animal models often examines only a single viral strain in a limited number of animals. We found that BG505’s ease of escape by single mutations is correlated with antibody breadth against natural sequences, but that there are also differences, especially among the broadest antibodies. We suggest that both measures may be useful for assessing the potential efficacy of bnAb-based therapies.

Of course, it is important to keep in mind that our maps only measure the effect single amino-acid mutations to BG505. Epistatic interactions among multiple mutations can play a role in viral fitness and immune evasion (Adams et al., 2017; Dahirel et al., 2011; Lynch et al., 2015b; Otsuka et al., 2018; Da Silva et al., 2010; Troyer et al., 2009; Wu et al., 2017). Similarly, we performed our experiments in the single genetic background of the BG505 Env, but the effects of mutations on viral growth and antigenicity sometimes differ among viral strains (Barton et al., 2016; Falkowska et al., 2014; Haddox et al., 2018).

Nonetheless, having complete antigenic maps of the BG505 Env versus a panel of important bnAbs provides a wealth of information that can help guide the study of HIV evolution and the development of anti-viral strategies. Future work that combines these antigenic maps with measurements (Haddox et al., 2016, 2018) or models (Louie et al., 2018; Shekhar et al., 2013; Zanini et al., 2017) of how mutations affect HIV’s replicative fitness could shed further light on the virus’s evolutionary dynamics under immune pressure.

## Methods

### Generation of mutant virus libraries

We have previously described the BG505.T332N mutant proviral DNA libraries and the resulting functional mutant virus libraries (Haddox et al., 2018). Briefly, triplicate mutant BG505.W6M.C2.T332N *env* libraries that contained randomized codon-level mutations to sites 31-702 (HXB2 numbering is used here and throughout this manuscript) were independently generated and cloned into Q23.BsmBI.ΔEnv proviral plasmid (Haddox et al., 2018). These proviral plasmid libraries, as well as wildtype proviral plasmid, were transfected into 293T cells (obtained from ATCC). We then passaged the transfection supernatant at an MOI of 0.01 TZM-bl infectious units/cell in SupT1.CCR5 cells. The resulting genotype-phenotype linked mutant virus libraries were concentrated via ultracentrifugation.

### Mutational antigenic profiling

The Env mutational antigenic profiling approach has been previously described (Dingens et al., 2017, 2018). Briefly, 0.5 - 1×10^6^ TZM-bl infectious units of independent mutant virus libraries were neutralized with each antibody at an ∼IC_95_ - IC_99.9_ concentration for one hour. Libraries were then infected into 1×10^6^ SupT1.CCR5 cells in R10 (RPMI, supplemented with 10% FBS, 1% 200 mM L-glutamine, and 1% of a solution of 10,000 units/mL penicillin and 10,000 µg/mL streptomycin) containing 100ug/mL DEAE-dextran. In parallel to each antibody selection, each mutant virus library was also infected into 1×10^6^ SupT1.CCR5 cells without antibody selection to serve as the experiment-specific non-selected control. Four 10-fold serial dilutions of each mutant virus library were also infected into 1×10^6^ cells as an infectivity standard curve. Cells were resuspended in 1 mL fresh R10 at three hours post infection. Cells were washed with PBS, pelleted, and non-integrated viral DNA was isolated via a miniprep at twelve hours post infection. The *env* gene was then amplified and sequenced using a barcoded subamplicon sequencing approach as previously described (Haddox et al., 2018), which introduces unique molecular identifiers to reduce sequencing error. The amount of viral genome in each sample was quantified via qPCR (Benki et al., 2006) or ddPCR (Dingens et al., 2017), and the fraction of each selected library that survived antibody selection relative to the non-selected control was interpolated from the infectivity standard curve.

### Analysis of deep sequencing data and computation of fraction surviving

We used dms_tools2 version 2.2.9 (https://jbloomlab.github.io/dms_tools2/) to analyze the deep sequencing data and calculate the fraction surviving (Bloom, 2015). The calculation of the fraction surviving statistic has been described previously (Doud et al., 2018) and is documented in detail at https://jbloomlab.github.io/dms_tools2/fracsurvive.html. Sequencing of wildtype proviral DNA plasmid was used as the error control during the calculation of the fraction surviving. We took the median values across all experimental replicates for each antibody, and plotted the excess fraction surviving data on logoplots rendered by dms_tools2 using weblogo (Crooks et al., 2004) and ggseqlogo (Wagih, 2017).

### Identification of significant sites of viral escape

Since the signal to noise ratio appeared to differ between antibodies, we defined statistically significant sites of viral escape beyond background individually for each antibody. For each antibody, we fit a gamma distribution to binned site fraction surviving values (median values across all replicates) using robust regression (soft L1 loss as implemented in scipy). We then identified sites that fell outside the range of values expected from this distribution at a false discovery rate of 0.01 (Figure S2). For the purposes of Figures 2B and 2C, multiple sites that disrupted a PNG that the antibody contacted was considered a single site of escape. However, when the antibody also contacted the second or third protein residues in the PNG and that site was site of significant escape, the site treated as additional site of escape.

### Structural analyses

Antibody contact sites were defined from Env-antibody structural models. The PDB models used were: 5FYK for VRC01 (Stewart-Jones et al., 2016), 5V8M for 3BNC117 (Lee et al., 2017), 5T3Z for 10-1074 (Gristick et al., 2016), 3U4E for PG9 (McLellan et al., 2011), 5V8L for PGT145 (Lee et al., 2017), 5FUU for PGT151 (Lee et al., 2016), 5I8H for VRC34.01 (Kong et al., 2016), 4G6F and for 10E8 (Huang et al., 2012). High resolution models of PGT121 bound to Env are not available; we used a model of PGT122 (PDB: 5FYL) (Stewart-Jones et al., 2016), which is a closely related “PGT121-like” clonal variant of PGT121 (Mouquet et al., 2012). Contact residues were defined as any Env residue where a non-hydrogen atom comes within 4 Å of any non-hydrogen antibody atom. When an antibody contacted a glycan, the N of that glycan’s PNG was counted as a contact. For asymmetric antibody-trimer structures, the closest distance of the three antibody-Env residue distances was used.

For Figures 3-5, we used the above referenced models (with one exception) to generate figures using PyMol. While we used the high resolution PG9-V2 scaffold structure (McLellan et al., 2011) to determine contact sites, we mapped the fraction surviving values onto the moderate resolution model of PG9 in complex with BG505 trimer (PDB: 5VJ6) (Wang et al., 2017) in Figure 3F to better illustrate the quaternary aspect of this apex epitope.

### TZM-bl neutralization assays

TZM-bl neutralization assays using BG505.T332N pseudoviruses bearing single additional point mutants were performed and analyzed as previously described (Dingens et al., 2017). The assay was performed in duplicate two or three independent times, and fold change in IC_50_ of each mutant relative to BG505.T332N wildtype pseudovirus was calculated independently for each experiment and then averaged across all replicates.

### Data availability and source code

Open-source software to analyze mutational antigenic profiling datasets is available at https://jbloomlab.github.io/dms_tools2/. The computational analysis is provided as an executable and HTML Jupyter notebook (File S2) and at https://github.com/jbloomlab/EnvsAntigenicAtlas, and the fraction surviving and excess fractions surviving values for each antibody are provided as File S3. Illumina deep sequencing reads are available from the NCBI SRA as study SRP157948, BioProject PRJNA486029, accession numbers SRX4553035-SRX4553088, SRX4614453.

## Acknowledgements

We thank Hugh Haddox and Shirleen Soh for contributing advice and code and Jeremy Roop and Caelan Radford for providing comments on this manuscript. We thank Rebecca Lynch, Ben Murrell, Katherine Bar, Craig Magaret, and Paul Edlefsen for providing insights into and viral sequences from bnAb immunotherapy clinical trials. We thank Peter Kwong for contributing VRC34.01. The following reagents were obtained through the NIH AIDS Reagent Program, Division of AIDS, NIAID, NIH: PG9 from Dr. Dennis Burton, 3BNC117 and 10-1074 from Dr. Michel Nussenzweig, PGT121 from Dr. Pascal Poignard, and 10E8 from Dr. Mark Connors.

ASD was supported by an NSF Graduate Research Fellowship (DGE-1256082). This work was supported by NIH grants R01AI127893 to JDB and by DA039543 and R01AI120961 to JO. JDB is an Investigator of the Howard Hughes Medical Institute.

## Author Contributions

Conceptualization, A.S.D.; Methodology, A.S.D., and J.D.B.; Validation, A.S.D., D.A., H.W.; Investigation, A.S.D.; Software, A.S.D. and J.D.B.; Supervision, J.O. and J.D.B.; Writing – Original Draft, A.S.D.; Writing – Review & Editing, A.S.D., J.O., and J.D.B.; Funding Acquisition, J.O. and J.D.B.

## Supplemental Figures and Files

**File S1 | The excess fraction surviving values plotted across the length of the mutagenized portion of Env for each antibody.** The underlay indicates contact sites, sites of significant escape, and the overlap between these groups of sites, as in Figure 2B, 2C.

**File S2 | The computational analysis.** A zip file containing an executable Jupyter notebook, an HTML version of the notebook, and all of the necessary input data to run the analysis.

**File S3 | The mutation fraction surviving and excess mutation fractions surviving datasets.** A zip file containing three CSV files for each antibody. One contains the mutation fraction surviving estimates, and one contains the excess mutation fraction surviving estimates plotted in the paper. The third contains *differential selection* estimates, which are log-transformed relative enrichment ratios that may be of use for certain purposes, such as examining mutations that are differentially depleted, rather than enriched, upon antibody selection. The differential selection metric is described in detail in (Doud et al., 2017) and at https://jbloomlab.github.io/dms_tools2/diffsel.html. Sites are numbered according to HXB2 reference strain numbering.

**Figure S1:**
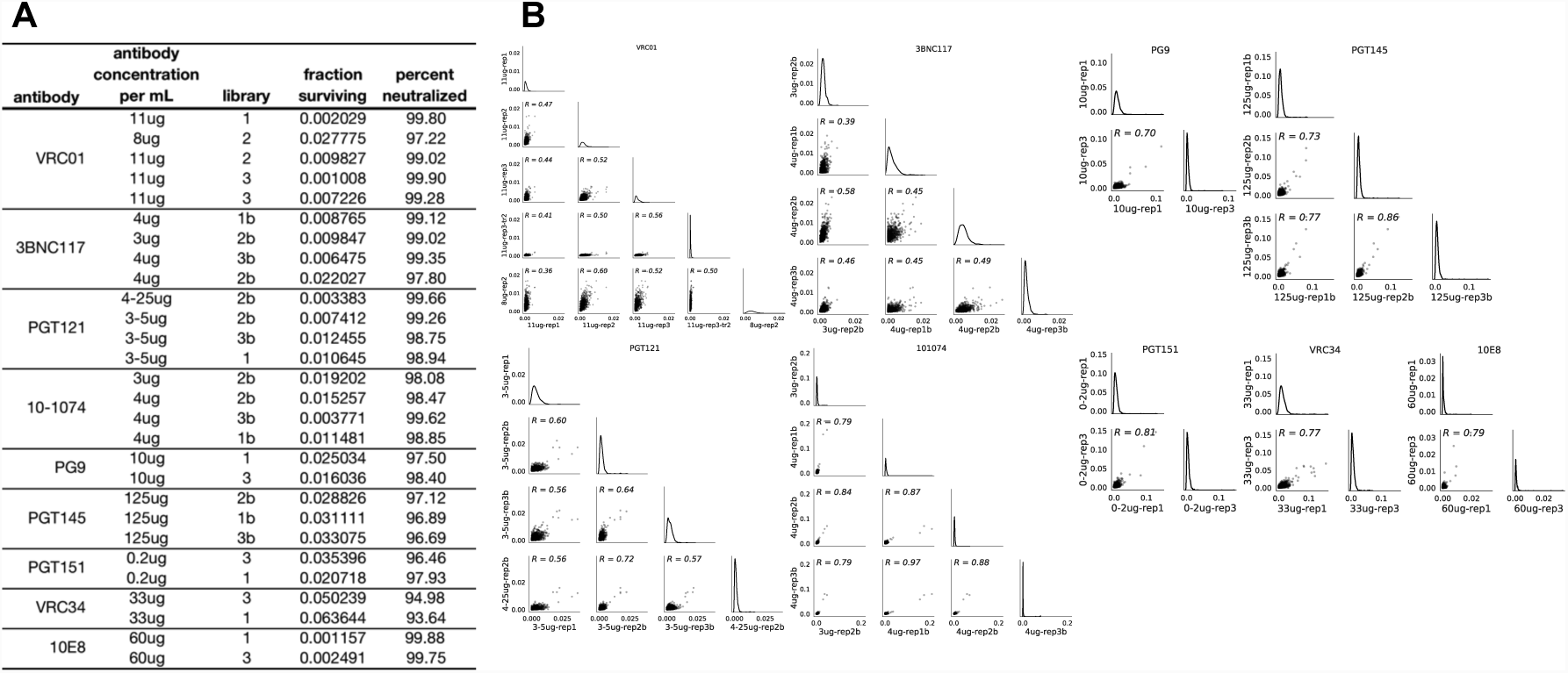
The fraction surviving measurements and correlation between mutational antigenic profiling biological replicates. Related to Figure 2. **A.** For each biological replicate, the antibody concentration used during the selection, which mutant virus library was used, and the fraction of that library that survived antibody selection is shown. For clarity, the percent neutralized (1-library fraction surviving) × 100 is also shown. **B.** The correlation between the average excess fraction surviving at each site for each biological replicate, for each antibody.

**Figure S2:**
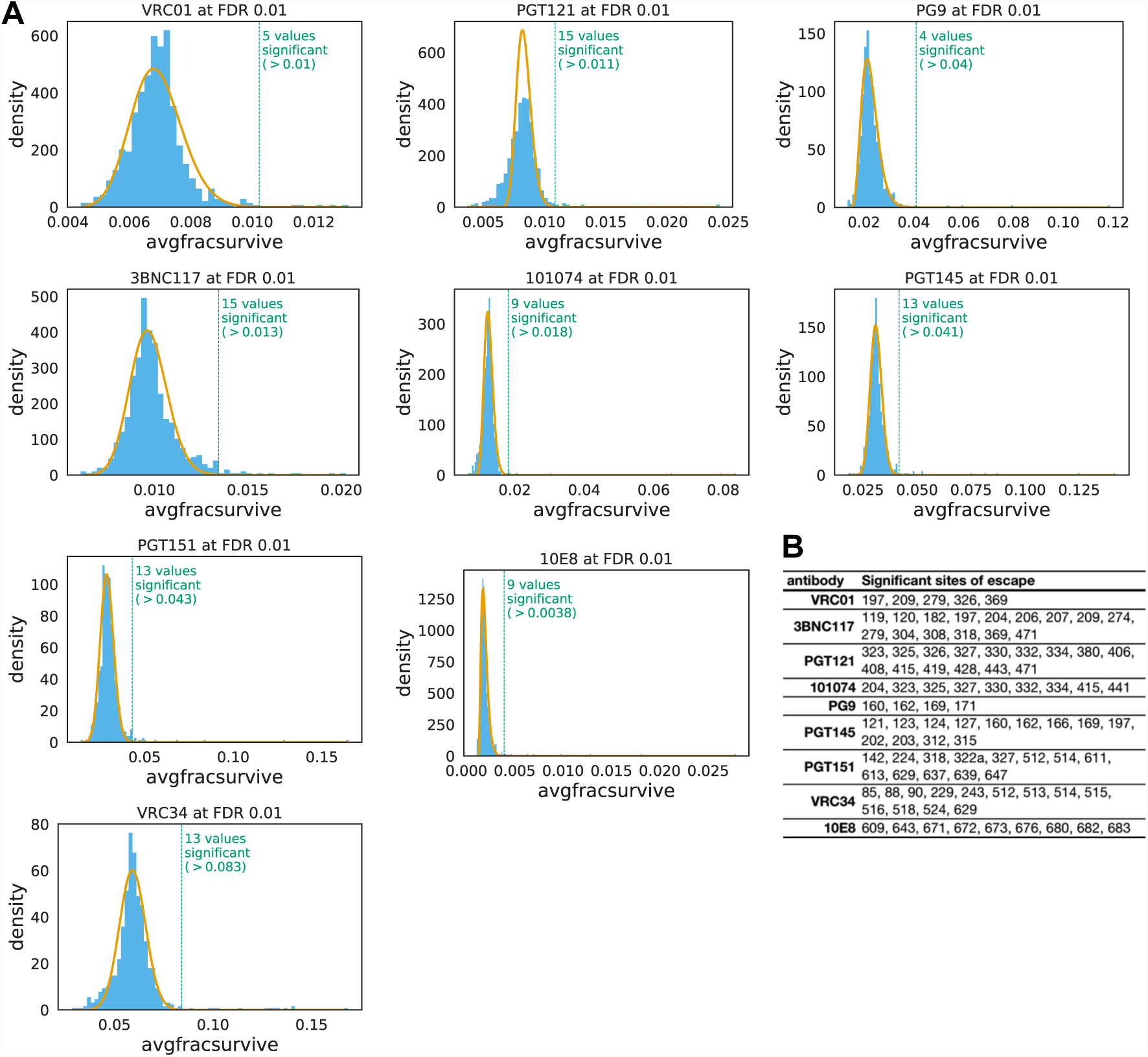
Identification of significant sites of viral escape. Related to Figure 2. **A.** For each antibody, the distribution of the average fraction surviving at each site is plotted in blue. A gamma distribution fit to the site fraction surviving values using robust regression is overlaid in yellow. A dotted line shows sites that fall beyond this distribution at a fall discovery rate of 0.01, and the number of sites that beyond this cutoff is labeled. Code that performs this analysis is at https://jbloomlab.github.io/dms_tools2/dms_tools2.plot.html#dms_tools2.plot.findSigSel. **B.** A table listing all of the significant sites of viral escape for each antibody.

**Figure S3:**
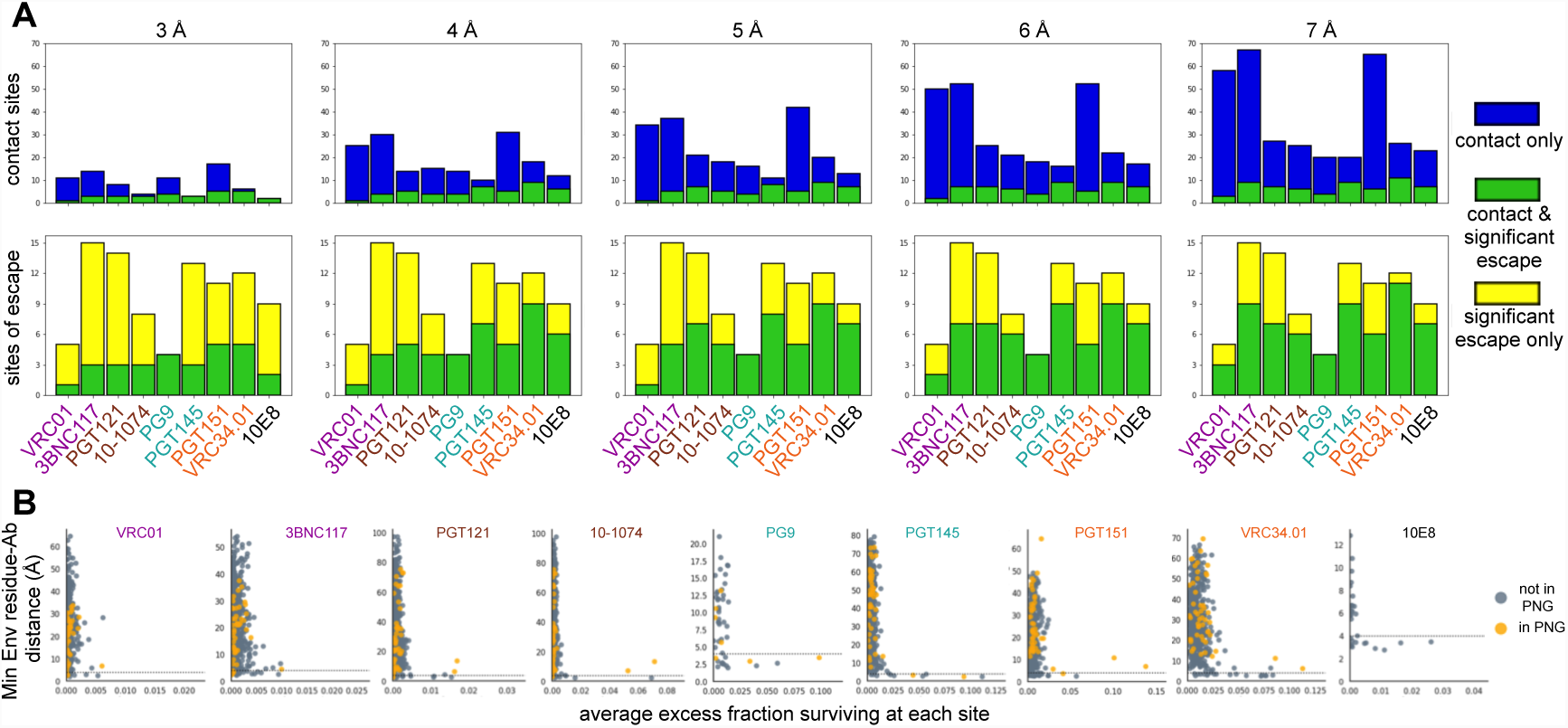
The overlap between each antibody’s structural and functional epitope. Related to Figure 2. **A.** As in Figure 2B and 2C, but using different distance cutoffs between non-hydrogen Env and antibody atoms to determine contact sites**. B.** For each Env residue in our library the minimum distance to the antibody is plotted against that site’s excess fraction surviving averaged across mutations. The 4 Å distance cutoff used in Figure 2B and 2C is plotted with a dotted line. Sites that fall within PNGs are colored yellow, while other sites are grey. Only sites that are in the structures used to determine distance cutoffs are plotted (see Methods for details of the structures used).

**Figure S4:**
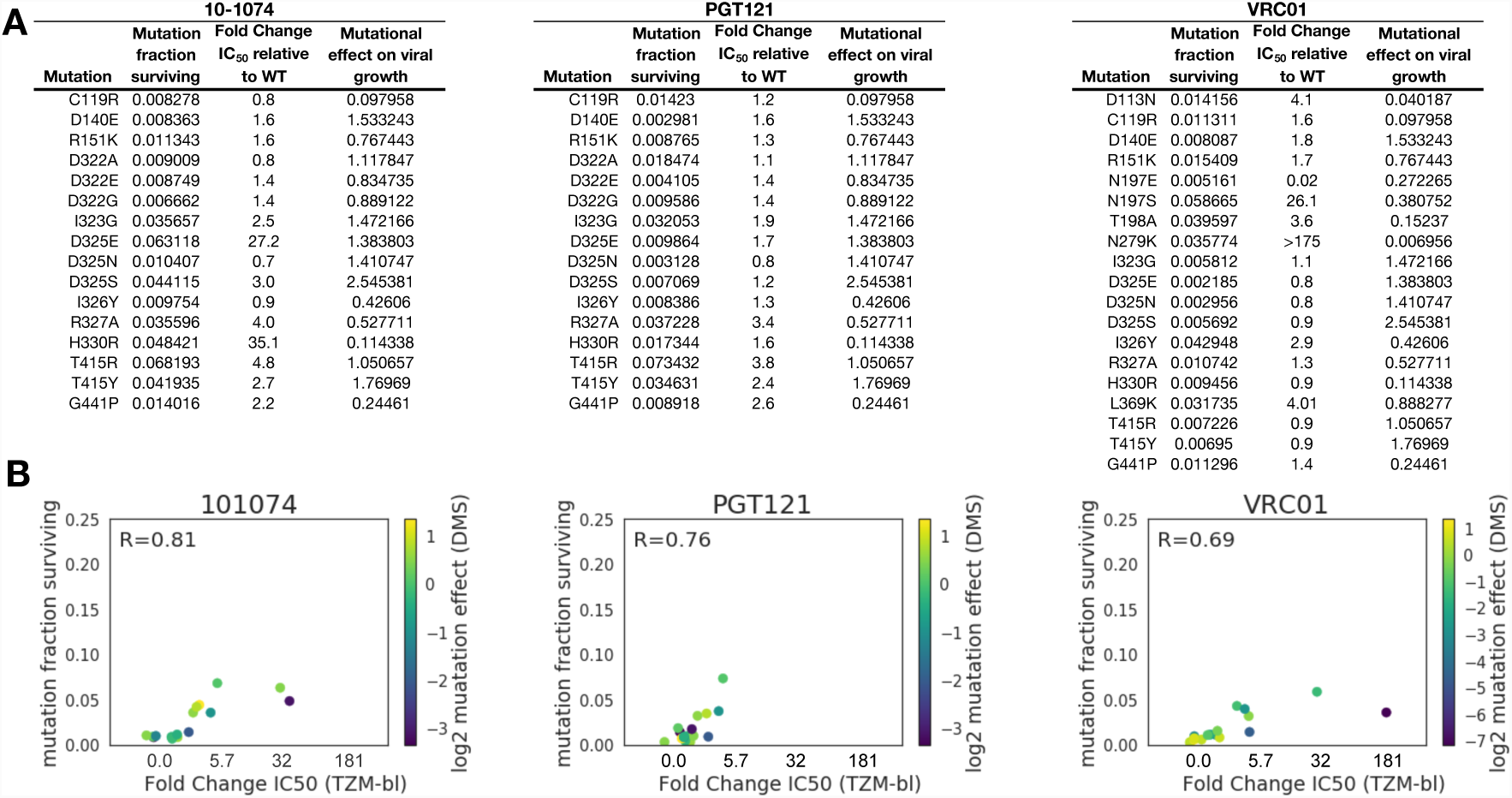
Validation of mutational antigenic profiling results using BG505 point mutants in TZM-bl neutralization assays. Related to Figure 2. **A.** The tables list mutations that were selected for validation in neutralization assays. For each mutation, the table gives the mutation’s fraction surviving antibody, fold change in IC_50_ in the TZM-bl neutralization assay, and the mutation’s effect on viral growth. The mutation’s effect on viral growth is calculated from our prior deep mutational scanning of Env for viral growth in cell culture, in the absence of any immune selection (Haddox et al., 2018). The mutational effect is the ratio of the preference for that mutant amino acid relative to the wildtype amino acid at that site. If the log mutational effect is >0, then that mutation grows better than wildtype in cell culture; if it is <0, that mutant grows worse than wildtype. **B.** For each antibody, we plot the correlation between the mutation fraction surviving and the fold change in IC_50_ relative to wildtype from TZM-bl assays. Points are colored according the log_2_ mutational effect of the mutation on viral growth. Where there are discrepancies between mutational antigenic profiling and TZM-bl assays, the mutational antigenic profiling estimate is usually less than would be predicted by the TZM-bl assay. These mutants (for example, N279K for VRC01 and H330R for 10-1074) often have a log_2_ mutational effects on viral growth <<0 (darker blue), indicating they are deleterious for viral growth in cell culture. Of note, mutating site 279 has also been shown to have a fitness cost in other assays (Lynch et al., 2015). Thus, the mutational antigenic profiling datasets may underestimate the antigenic effect of high fitness cost mutations, though these measures may possibly better reflect the ability of a mutant to both replicate and escape antibody neutralization *in vivo* than TZM-bl point mutant neutralization sensitivities alone.

**Figure S5:**
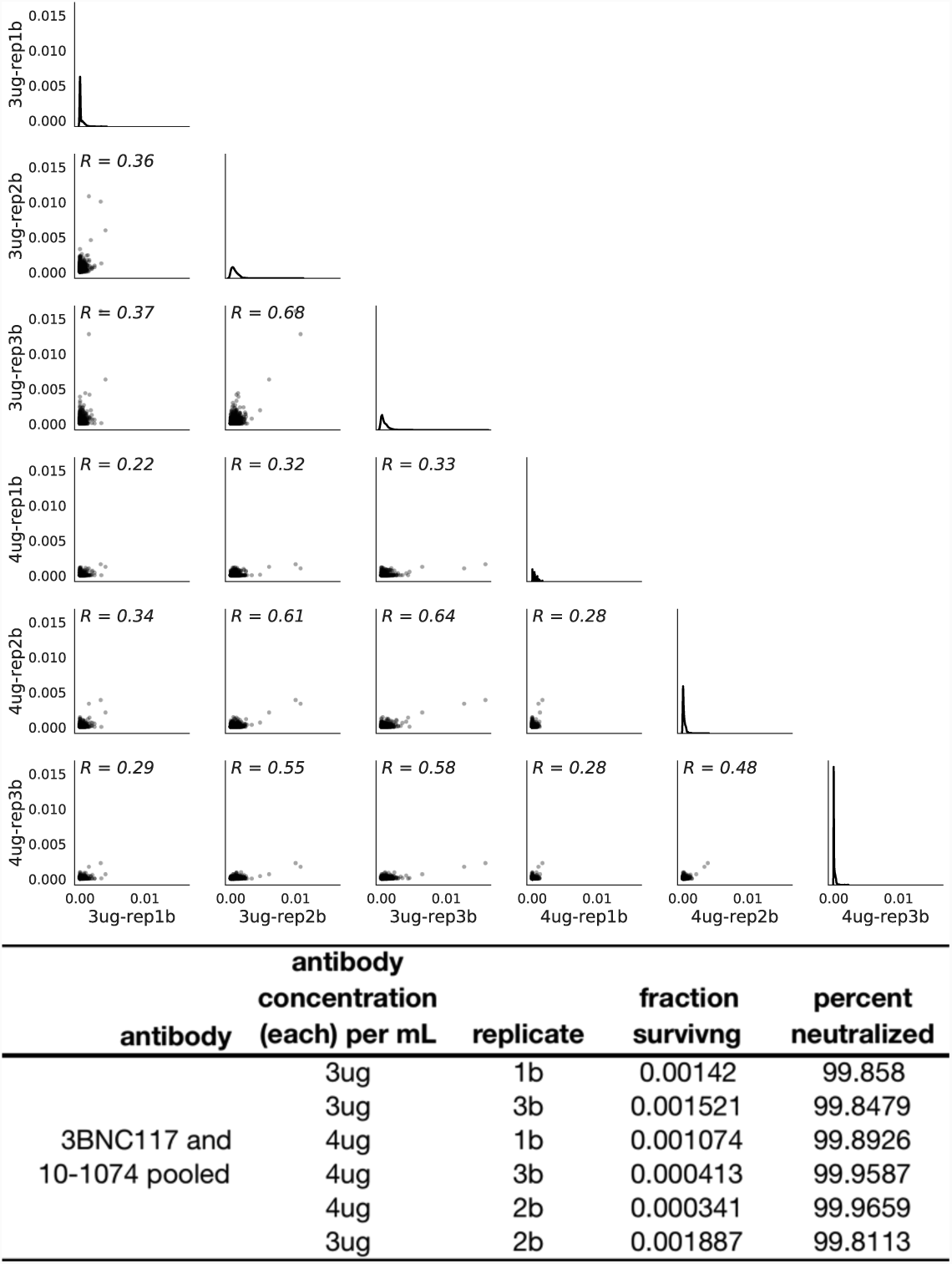
The correlation between mutational antigenic profiling biological replicates for the pooled 3BNC117 and 10-1074 antibodies. Related to Figure 6. **A**. The correlation between the excess fraction surviving averaged across mutations at each site for each biological replicate of the pooled antibody selections. **B.** For each biological replicate, the antibody concentration used during the selection, which mutant virus library was used, and the fraction of that library that survived antibody selection is shown. For clarity, the percent neutralized (1-library fraction surviving) × 100 is also shown.

**Table S1:**
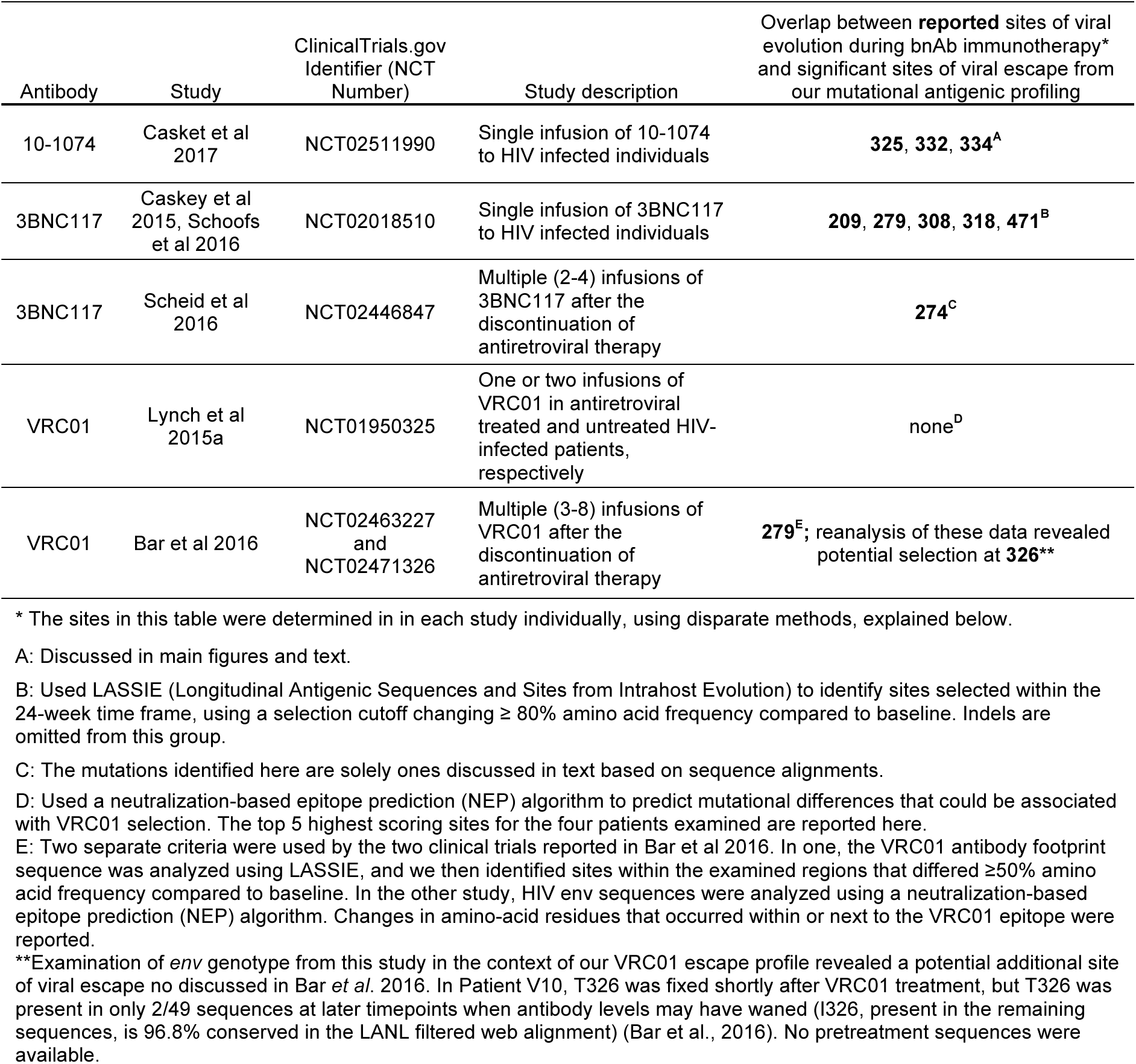
Overlap between mutational antigenic profiling sites of escape and sites of viral evolution that occurred *in vivo* during bnAb immunotherapy. Related to Figure 3 and 4.

| Antibody | Study | ClinicalTrials.gov Identifier (NCT Number) | Study description | Overlap between **reported** sites of viral evolution during bnAb immunotherapy* and significant sites of viral escape from our mutational antigenic profiling |
| --- | --- | --- | --- | --- |
| 10-1074 | Casket et al 2017 | NCT02511990 | Single infusion of 10-1074 to HIV infected individuals | **325, 332, 334<sup>A</sup>** |
| 3BNC117 | Caskey et al 2015, Schoofs et al 2016 | NCT02018510 | Single infusion of 3BNC117 to HIV infected individuals | **209, 279, 308, 318, 471<sup>B</sup>** |
| 3BNC117 | Scheid et al 2016 | NCT02446847 | Multiple (2-4) infusions of 3BNC117 after the discontinuation of antiretroviral therapy | **274<sup>C</sup>** |
| VRC01 | Lynch et al 2015a | NCT01950325 | One or two infusions of VRC01 in antiretroviral treated and untreated HIV-infected patients, respectively | none<sup>D</sup> |
| VRC01 | Bar et al 2016 | NCT02463227 and NCT02471326 | Multiple (3-8) infusions of VRC01 after the discontinuation of antiretroviral therapy | **279<sup>E</sup>**; reanalysis of these data revealed potential selection at **326<sup>\*\*</sup>** |

\* The sites in this table were determined in in each study individually, using disparate methods, explained below.
A: Discussed in main figures and text.
B: Used LASSIE (Longitudinal Antigenic Sequences and Sites from Intrahost Evolution) to identify sites selected within the 24-week time frame, using a selection cutoff changing $\geq 80\%$ amino acid frequency compared to baseline. Indels are omitted from this group.
C: The mutations identified here are solely ones discussed in text based on sequence alignments.
D: Used a neutralization-based epitope prediction (NEP) algorithm to predict mutational differences that could be associated with VRC01 selection. The top 5 highest scoring sites for the four patients examined are reported here.
E: Two separate criteria were used by the two clinical trials reported in Bar et al 2016. In one, the VRC01 antibody footprint sequence was analyzed using LASSIE, and we then identified sites within the examined regions that differed $\geq 50\%$ amino acid frequency compared to baseline. In the other study, HIV env sequences were analyzed using a neutralization-based epitope prediction (NEP) algorithm. Changes in amino-acid residues that occurred within or next to the VRC01 epitope were reported.
**\*\***Examination of *env* genotype from this study in the context of our VRC01 escape profile revealed a potential additional site of viral escape not discussed in Bar et al. 2016. In Patient V10, T326 was fixed shortly after VRC01 treatment, but T326 was present in only 2/49 sequences at later timepoints when antibody levels may have waned (I326, present in the remaining sequences, is 96.8% conserved in the LANL filtered web alignment) (Bar et al., 2016). No pretreatment sequences were available.

